# *In silico* prediction of blood cholesterol levels from genotype data

**DOI:** 10.1101/503003

**Authors:** Francesco Reggiani, Marco Carraro, Anna Belligoli, Marta Sanna, Chiara dal Prà, Francesca Favaretto, Carlo Ferrari, Roberto Vettor, Silvio C.E. Tosatto

## Abstract

In this work we present a framework for blood cholesterol levels prediction from genotype data. The predictor is based on an algorithm for cholesterol metabolism simulation available in literature, implemented and optimized by our group in R language. Main weakness of the former simulation algorithm was the need of experimental data to simulate mutations in genes altering the cholesterol metabolism. This caveat strongly limited the application of the model in the clinical practice. In this work we present how this limitation could be bypassed thanks to an optimization of model parameters based on patients cholesterol levels retrieved from literature. Prediction performance has been assessed taking in consideration several scoring indices currently used for performance evaluation of machine learning methods. Our assessment shows how the optimization phase improved model performance, compared to the original version available in literature.

## Introduction

Recent exome-wide association studies [1] started to shed light on the complex genomic architecture behind the regulation of blood cholesterol levels in humans. Reliable tools to predict human cholesterol levels from genotype are not available yet. The huge number of genes involved in the regulation of this trait and the complex interaction with environmental factors as diet, sex and age make modelling cholesterol levels a difficult task. However, particular situations exist where a single mutation is related to significant variations of cholesterol levels. Example are damaging mutations on genes involved in hepatic uptake of Low Density Lipoprotein (LDL), as the Low Density Lipoprotein Receptor (LDLR) gene, causing familial hypercholesterolemia characterized by elevated levels of LDL and total plasma cholesterol but with normal concentrations of triglycerides [2]. Other processes involved in cholesterol metabolism are affected by genetic mutations, with a wide range of phenotypes depending on the gene involved, like marked High Density Lipoprotein (HDL) cholesterol levels deficiency as seen in patients affected by Tangier disease [3]. The aim of this work is to test the reliability of a modelling approach aimed to predict cholesterol levels relying on patient’s genotype data only. Effective way to simulate in silico metabolism are dynamic models. In this kind of simulations, the development of the system in time is computed through a set of ordinary differential equations, able to simulate the variations of chemical species concentration. Several information are required for the development of these models: interactions between the chemical species involved in the biological process, kinetic parameters associated to chemical reactions occurring in the system and its initial state. The simulation of a biological perturbation could be obtained by modifying model parameters (e.g. decreasing kinetic rates) and observing variations occurring in the system [4]. Several *in silico* models simulating cholesterol metabolism have been proposed so far, both for human and animal models [5]. In this work we started from an algorithm for cholesterol metabolism simulation published in literature by van de Pas and colleagues in 2012 [6]. This physiologically based kinetic model is based on differential equations, computing the flow of cholesterol in different body organs. The whole process is regulated by a set of rates, each one related to a gene that has a key role in cholesterol metabolism. Simulation of mutations effects depends on reducing rates (*f_mut_*) estimated from wet lab experiments. This kind of information is usually not easily accessible, strongly limiting the usability of the model. In this work we implemented and optimized the framework for blood cholesterol levels prediction making it able to perform reliable predictions when only patient’s genotype data are available. The model has been improved through a training phase, in which reducing rates (*f_mut_*) were estimated from phenotype data of patients affected by mutations on key regulatory genes of cholesterol metabolism. Assessment measures confirmed how the optimized model presents improved performance, reducing the error between experimental and predicted data, compared to the original version available in literature [6].

## Materials and methods

### *In silico* kinetic model for cholesterol levels prediction

An available in silico kinetic model [6] has been used as basis for predicting plasma cholesterol concentrations in humans. The kinetic model was developed to simulate cholesterol levels for a reference man of 70 kg. The model is composed of 8 pools, representing main sites of cholesterol storage in the human body (Fig 1). These pools can be grouped in 4 main entities corresponding to plasma, intestine, liver and periphery. Each cholesterol pool is modeled by a differential equation, composed by a set of rates moving cholesterol from or to a different one. These pools are connected by 21 kinetic rates, each one representing the main gene responsible of regulating that specific biochemical reaction (Table 1).

**Fig 1.**
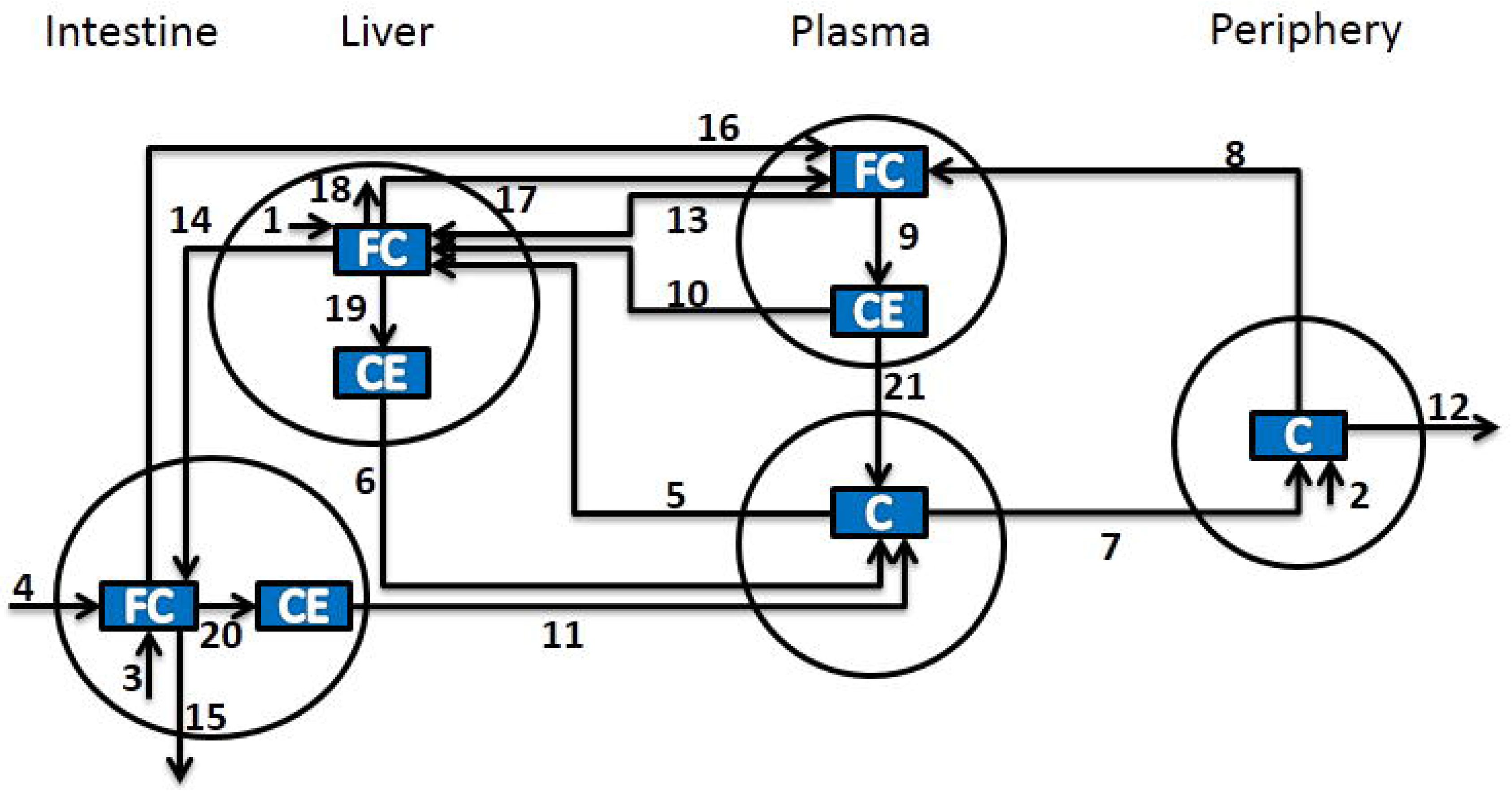
Conceptual model for pathways and genes determining cholesterol plasma levels used van de Pas and colleagues. [6], [7]. Process numbers stand for: 1, hepatic cholesterol synthesis (DHCR7); 2, peripheral cholesterol synthesis(DHCR7); 3, intestinal cholesterol synthesis (DHCR7); 4, dietary cholesterol intake (NPCIL1); 5, hepatic uptake of cholesterol from LDL (LDLR,APOB,APOE); 6, VLDL-C secretion (MTTP); 7, peripheral uptake of cholesterol from LDL (LDLR,APOB,APOE); 8, peripheral cholesterol transport to HDL (ABCA1); 9, HDL-associated cholesterol esterification (LCAT); 10, hepatic HDL-CE uptake (SCARB1); 11, intestinal chylomicron cholesterol secretion (MTTP); 12, peripheral cholesterol loss; 13, hepatic HDL-FC uptake (MTTP); 14, biliary cholesterol excretion (ABCG8,NPC1L1); 15, fecal cholesterol excretion; 16, intestinal cholesterol transport to HDL (ABCA1); 17, hepatic cholesterol transport to HDL (ABCA1); 18, hepatic cholesterol catabolism (CYP7A1); 19, hepatic cholesterol esterification (SOAT2); 20, intestinal cholesterol esterification (SOAT2); and 21, CE transfer from HDL to LDL (CETP).

**Table 1.**
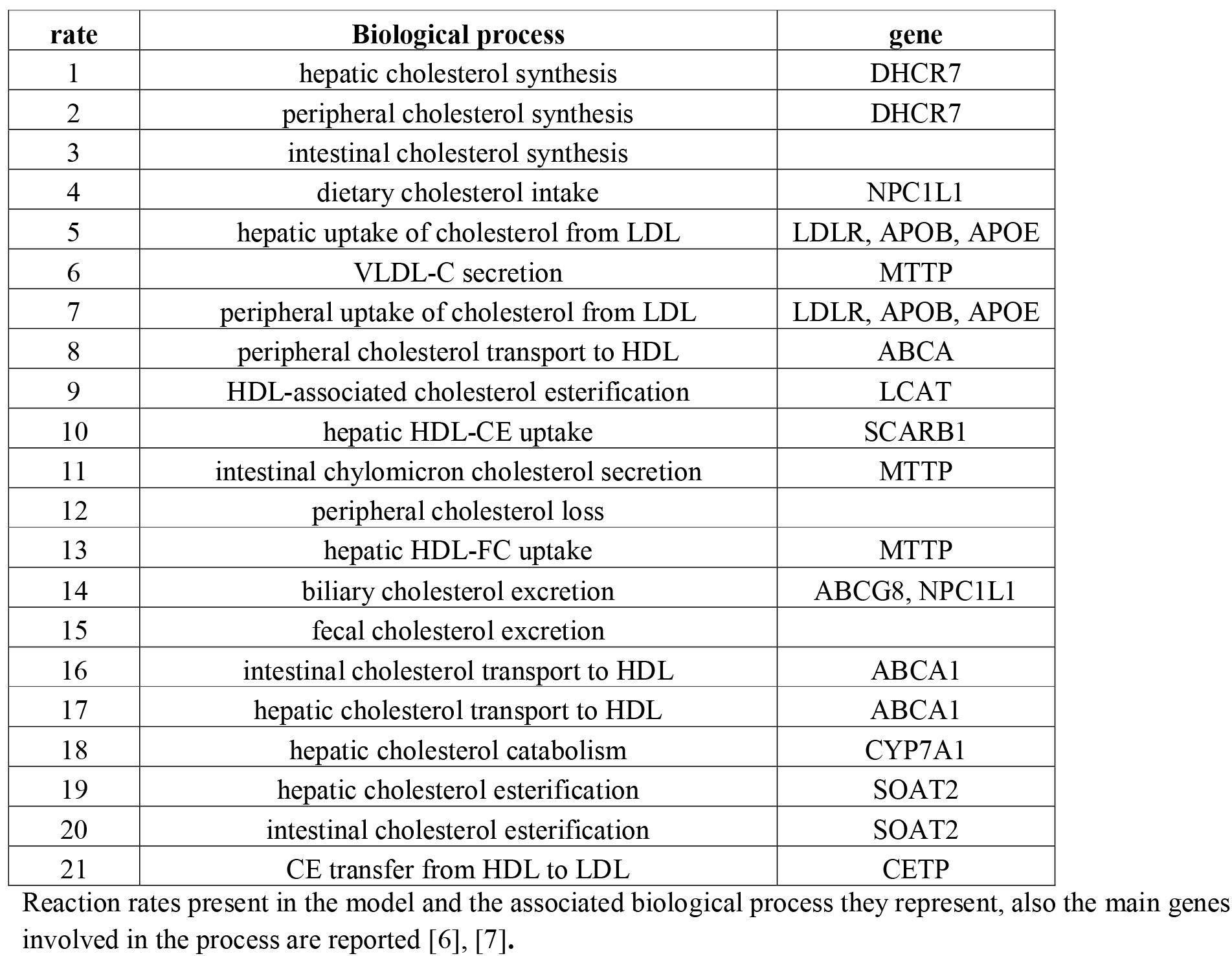
Biological process and genes associated to each rate of the model.

Rates depend on kinetic constants, organ volumes, body weight and pool cholesterol concentrations. In the original model, all parameters have been computed from data published in literature [6]. The model was calibrated to immediately reach a steady state, a stable equilibrium in which each compartment has a constant cholesterol concentration in time. To simulate a mutation affecting the activity of a gene, a set of rate reduction parameters (*f_mut_*), each one in the interval [0, 1], multiplies the standard rates to represent the effect of the mutated genes. These values were computed on the basis of experimental data available in literature. Example is the value of the *f_mut_* related to mutations affecting the gene CYP7A1 involved in byle acid synthesis, where the rate reduction parameter was computed as the ratio of bile acids contents in the stools of patients carring the mutation over controls [6].

These kind of perturbations force a re-tuning of the system, moving from the original steady state to a new one, where blood cholesterol profiles were comparable to the real values detected in patients affected by that particular mutation.

### Model implementation

The algorithm of the available physiologically based kinetic mode [5], was implemented in R language [8].

The *deSolve* package [9] was used for solving differential equations.

New *f_mut_* values have been obtained thanks to a training procedure exploiting a dataset composed of cholesterol levels and genotypes of mutated patients. This operation required the usage of the Levenberg-Marquardt algorithm as implemented in the *Minpack.lm* package [10].

### Training phase

To improve performance in predicting genetic mutations’ effect on cholesterol levels, *f_mut_* parameters, each one related to a particular gene mutation and rates of the model, had been trained on phenotype data of a custom dataset of patients. The Levenberg-Marquardt minimization method has been used to estimate the *f_mut_* parameters able to minimize the difference between predicted and experimentally measured level of HDL and LDL, divided by the control, intended as level of cholesterol of the model when no mutation is present (Equation 1 and 2). Exceptions are patients affected by mutations on the DHCR7 genes where only total cholesterol (TC) levels were found in literature. In this case the difference between real and predicted total cholesterol rate where taken in account (Equation 3).

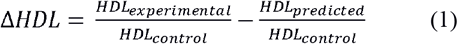

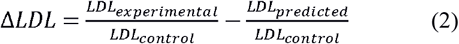

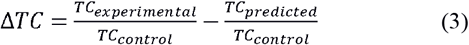

The optimized *f_mut_* parameters are reported in Table 2.

**Table 2.**
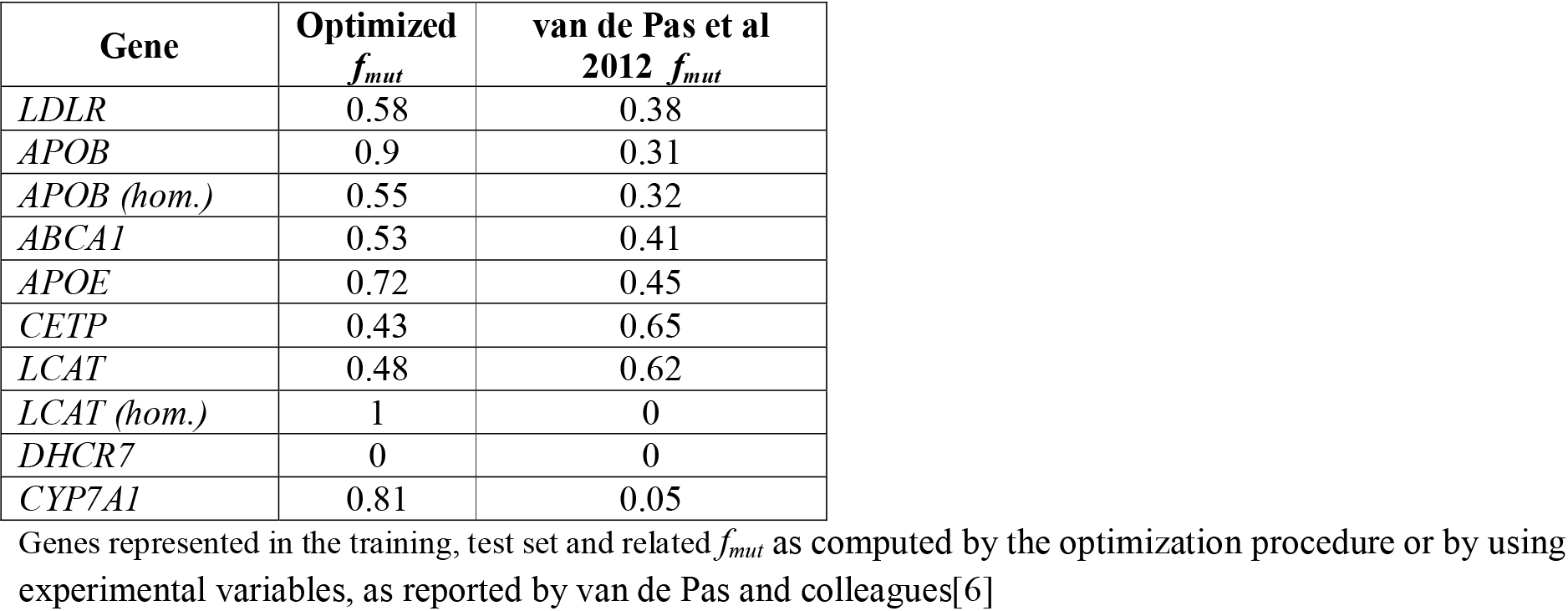
Optimized *f_mut_* parameters and related genes.

| Gene | Optimized $f_{mut}$ | van de Pas et al 2012 $f_{mut}$ |
| --- | --- | --- |
| *LDLR* | 0.58 | 0.38 |
| *APOB* | 0.9 | 0.31 |
| *APOB (hom.)* | 0.55 | 0.32 |
| *ABCA1* | 0.53 | 0.41 |
| *APOE* | 0.72 | 0.45 |
| *CETP* | 0.43 | 0.65 |
| *LCAT* | 0.48 | 0.62 |
| *LCAT (hom.)* | 1 | 0 |
| *DHCR7* | 0 | 0 |
| *CYP7A1* | 0.81 | 0.05 |

Genes represented in the training, test set and related $f_{mut}$ as computed by the optimization procedure or by using experimental variables, as reported by van de Pas and colleagues[6]

### Training set

The training set is represented by a custom database of patients affected by single mutations (either in homozygous or heterozygous form), in one of the key genes regulating cholesterol metabolism (Fig 2). For each patient the levels of HDL, LDL or total cholesterol and the causative mutation were extracted from literature. Each gene is covered by a different number of individuals due to the relative abundance of works in literature (Table 3). Special cases are the CETP gene, where only information regarding the mean levels of blood cholesterol were found in literature and the DHCR7 gene, where only the levels of blood total cholesterol were found. The training set has been divided in two sections (Table 3). The first group is represented by hypercholesterolemic patients with mutations affecting a set of genes involved in the development of Autosomal Dominant Hypercholesterolemia [11]: LDLR, APOB and APOE genes, represented by reaction 5 and 7 of the model (Table 1). The second part of the dataset is composed of patients with damaging mutations on 5 different genes: ABCA1, CETP, LCAT, DHCR7 and CYP7A1 (affected rates are shown in Table 1). Patients of the Autosomal Dominant Hypercholesterolemia dataset are characterized by high levels of LDL cholesterol, while the second part of the dataset is composed by different ranges of HDL and LDL, depending on the gene affected by the mutation (Figure 2).

**Fig 2.**
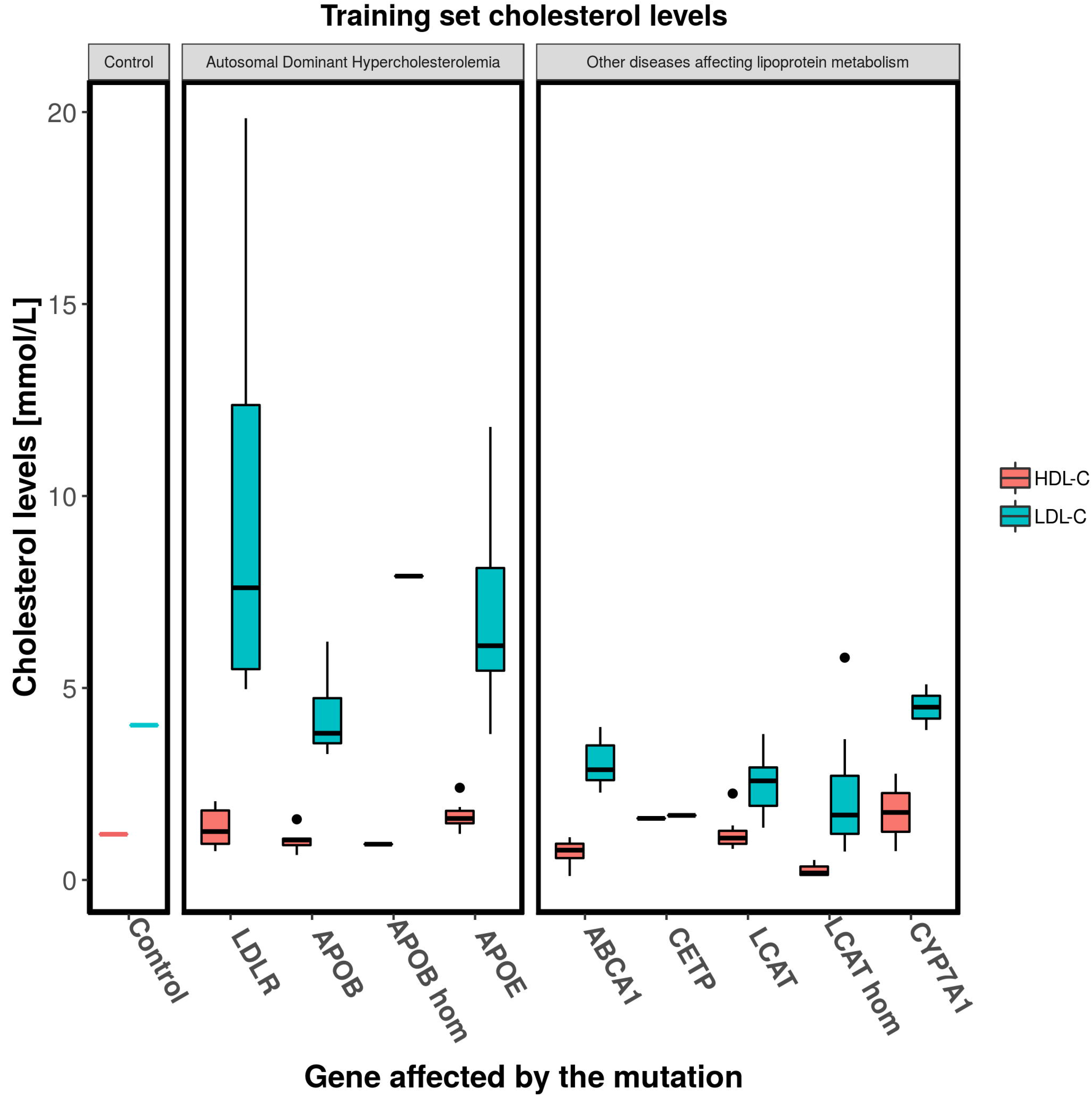
Training set patients cholesterol levels. Boxplot of HDL and LDL cholesterol levels of the patients composing the training set. From left to right: cholesterol levels of the model at the steady state, patients affected by Autosomal Dominant Hypercholesterolemia (with high levels of LDL and low HDL), patients affected by other disease altering lipoprotein metabolism.

**Table 3.**
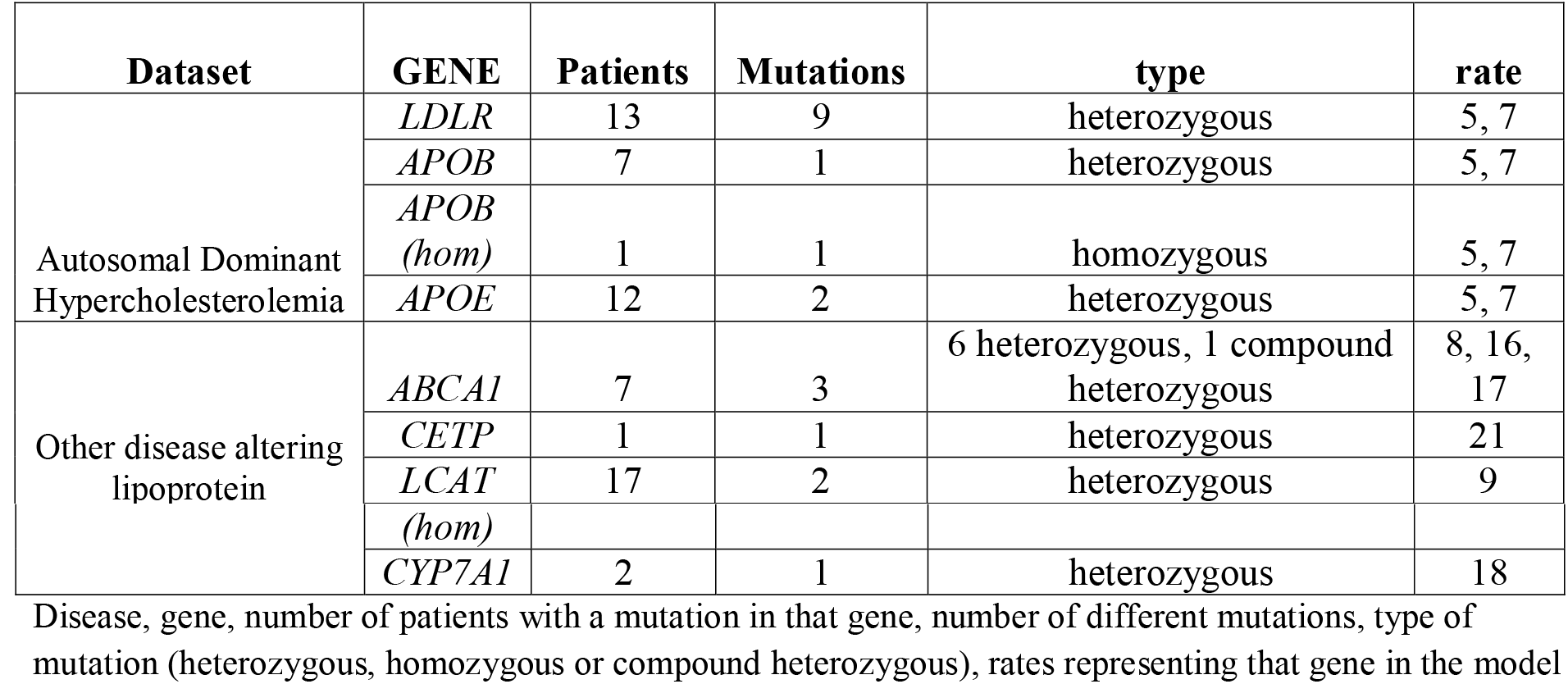
Training set composition.

### Test phase

Prediction performance were tested on a dataset retrieved form literature [6]. The dataset is the same one used to test performance of the former version of the model. This test set has been used in order to highlight performance comparison between the versions of the algorithm. The effect of a genetic mutation was simulated for each individual of the dataset until a steady state was reached (fixed threshold: 1000 days). Predicted HDL, LDL and total cholesterol were than compared to experimental data.

### Test set

Test set is composed by patients affected by 10 mutations. All mutations affects genes present in the training set of this work. The first group of mutations maps on the LDLR, APOB and APOE genes, involved in hepatic cholesterol uptake. Patients affected by this kind of mutations have high levels of LDL and total cholesterol. Genetic mutations affecting the other genes of the dataset have different effects on lipid profiles. Mutations on the ABCA1 can cause marked HDL cholesterol levels deficiency as reported for different disease like hypoalphaliproproteinemia or Tangier disease [3]. CETP is a protein involved in the transport of cholesterol esters from HDL to LDL, deficiency of this protein can cause a marked increase of HDL levels [3]. LCAT is a gene involved in cholesterol esterification in HDL particles, mutation on this gene can cause LCAT deficiency, characterized by low levels of HDL and LDL cholesterol [3]. Patients with mutations in heterozygous or homozygous form has been included in the training set. DHCR7 gene is responsible for the last step of the cholesterol biosynthesis pathway. Reduced enzyme activity cause low levels of blood cholesterol, as reported in patients affected by the Smith-Lemli-Opitz syndrome [12]. CYP7A1 gene is involved in cholesterol catabolism and bile acids synthesis, mutations affecting this gene cause an increase of total, hepatic cholesterol and a decrease in bile acids secretion [13].

### Performance assessment

The assessment approach used in this work was influenced by the methods used for the evaluation of tools predicting the effect of variants on continuous phenotypes [14]. Model performance has been evaluated in terms of distance and correlation, measuring the deviation from experimental values while assessing model capability to predict a decrease or increase of cholesterol levels. The analysis has been conducted at two levels. In the first part of the assessment, predictions were evaluated at the level of the single gene in order to understand if prediction error was homogeneous or significantly different for some of the mutations. The second part of the assessment focused on the overall performance of the predictor. In the first phase, the analysis was focused on assessing model performance in terms of prediction error computed on each element of the test set: the deviation was evaluated by computing the difference between predicted and experimental data, in terms of the rate of cholesterol levels in case and control. This analysis was aimed to highlight mutation effects that where under or over-predicted. In the second part of the assessment, model performance has been evaluated in terms of correlation and error measures on the whole dataset. Correlation measures used for the assessment has been: Pearson and Kendall’s tau correlation coefficients. To better understand the amount of variability described by the model compared to the variability inside the data, the R^2^ index was used. RMSE (Root Mean Squared Error) has been used in order to evaluate if the method predicting cholesterol levels with huge deviation from real ones. A sensitivity analysis was performed on a set of rates, corresponding to genes represented in the test set. The aim of this analysis was to understand the effect of a perturbation of specific model parameters on the output [4]. In this case we decreased rates associated to genes represented in the test set, using a reducing factor [0.1, 1] and measured model cholesterol levels when a steady state was reached.

## Results and discussion

### Performance assessment on single genes mutations

The first part of the assessment was aimed to understand how the model performs on the single mutations represented in the test set. This type of analysis highlighted cases where the model overestimated or underestimated cholesterol levels, respectively called positive or negative errors. The error represent the increase or decrease of cholesterol in case relative to controls (equation 1, 2, 3), which is not observed in experimental data. The errors where divided by real data and converted to percentages as reported in Table 4. As already introduced, the datasets of patients have been divided in two sections. The first group of elements of the test set is composed by patients affected by damaging mutations on genes that have a role in the onset of the Autosomal Dominant Hypercholesterolemia: LDLR, APOB and APOE. The main effect of simulating these mutations is an increase of blood cholesterol levels of LDL and decrease of HDL (Table 5), as observed in real cases [15]. The algorithm predicted cholesterol levels caused by mutations in LDLR and APOB with a reduced error intervals: [−35.3%, 11.5%] for HDL, [−26.2%, −12.9%] for LDL and [−20.5%, – 13.7%] for total cholesterol, respect to the former version of the model. The original model in fact, shown to drive predictions towards an overestimation of the mutation effect, as shown by the prediction errors of HDL [−52%, −30.8%], LDL [35%, 139.7%], and total cholesterol [29.7%, 115.9%]. A particular cases is the one of mutations in APOE, where the algorithm strongly underestimated the effect of damaging mutations on total cholesterol levels. In this case a higher error has been registered for our optimized model (−53.1%) compared with the former version (−27.4%). This situation is mainly related to the fact that the average levels of total cholesterol of patients in the training set was lower than the one of the test set.

**Table 4.** Models predictions percentage error on elements of the test set.

| Mutation | Gene | HDL vdP et al | LDL vdP et al | TC vdP et al | HDL train | LDL train | TC train |
| --- | --- | --- | --- | --- | --- | --- | --- |
| 1 | LDLR | -30.83 | 35.05 | 29.65 | -13.52 | -14.82 | -13.69 |
| 2 | APOB | -36.7 | 139.71 | 115.91 | 11.53 | -26.23 | -20.45 |
| 3 | APOB (hom) | -51.96 | 62.66 | 49.05 | -35.34 | -12.89 | -15.14 |
| 4 | ABCA1 | 147.99 | -57.44 | -44.77 | 200.05 | -51.25 | -35.99 |
| 5 | APOE | NA | NA | -27.4 | NA | NA | -53.15 |
| 6 | CETP | 12.7 | -4.82 | -0.72 | 34.04 | -11.53 | -0.46 |
| 7 | LCAT | 34.3 | -0.66 | 21.7 | 39.18 | -3.08 | 20.55 |
| 8 | LCAT (hom) | 679.37 | -19.37 | 10.11 | 426.32 | 21.95 | 29.87 |
| 9 | DHCR7 | NA | NA | 171.51 | NA | NA | 171.51 |
| 10 | CYP7A1 | -4.42 | -42.09 | -34.15 | 1.37 | -50.07 | -40.81 |

Mutation numeric ID, gene, HDL, LDL and total cholesterol error (as percentage of experimental value), of predictions based on $f_{mut}$ as reported by van de Pas and colleagues[6], or trained $f_{mut}$ .

**Table 5.**
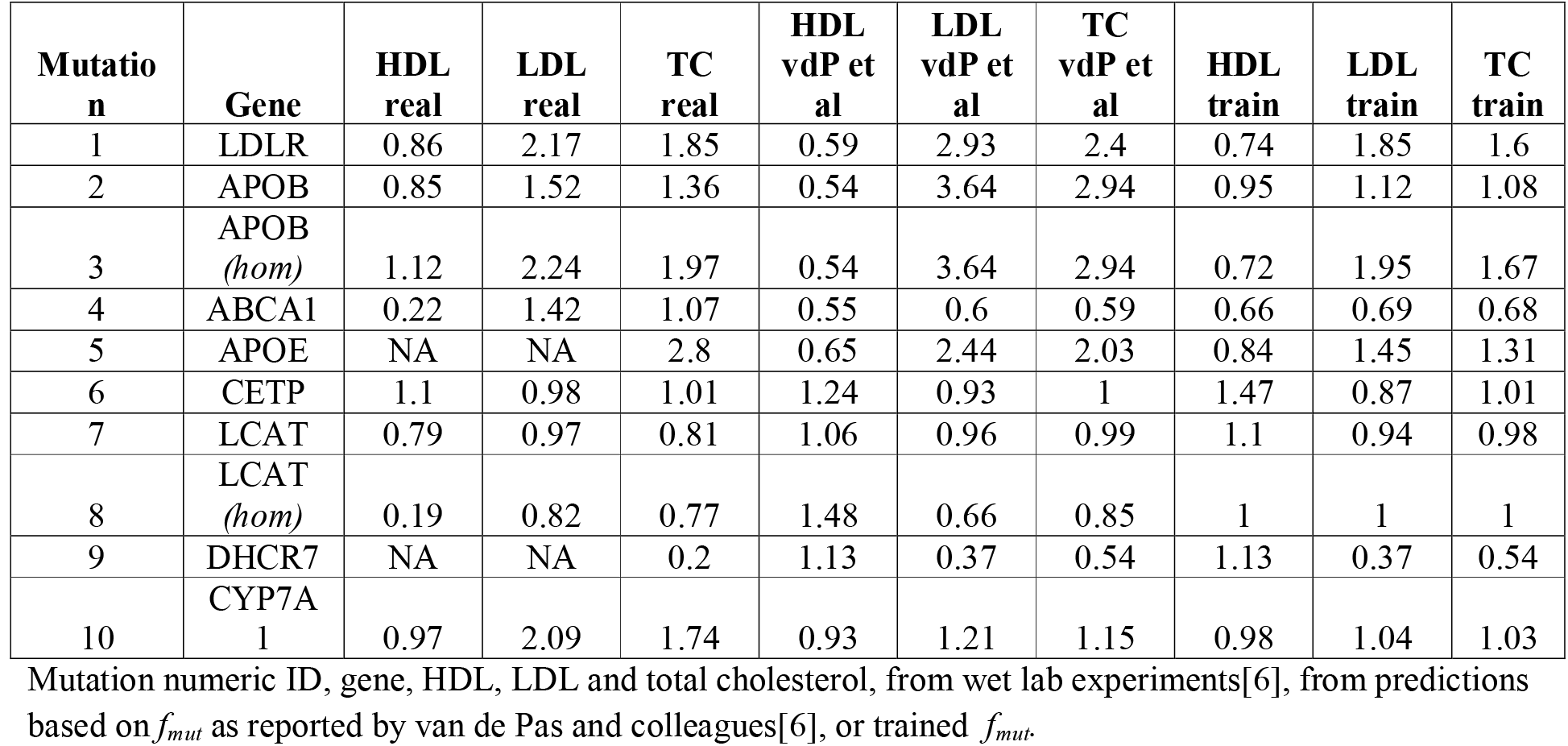
Experimental and predicted cholesterol levels of the blind set.

| Mutation | Gene | HDL real | LDL real | TC real | HDL vdP et al | LDL vdP et al | TC vdP et al | HDL train | LDL train | TC train |
| --- | --- | --- | --- | --- | --- | --- | --- | --- | --- | --- |
| 1 | LDLR | 0.86 | 2.17 | 1.85 | 0.59 | 2.93 | 2.4 | 0.74 | 1.85 | 1.6 |
| 2 | APOB | 0.85 | 1.52 | 1.36 | 0.54 | 3.64 | 2.94 | 0.95 | 1.12 | 1.08 |
| 3 | APOB (hom) | 1.12 | 2.24 | 1.97 | 0.54 | 3.64 | 2.94 | 0.72 | 1.95 | 1.67 |
| 4 | ABCA1 | 0.22 | 1.42 | 1.07 | 0.55 | 0.6 | 0.59 | 0.66 | 0.69 | 0.68 |
| 5 | APOE | NA | NA | 2.8 | 0.65 | 2.44 | 2.03 | 0.84 | 1.45 | 1.31 |
| 6 | CETP | 1.1 | 0.98 | 1.01 | 1.24 | 0.93 | 1 | 1.47 | 0.87 | 1.01 |
| 7 | LCAT | 0.79 | 0.97 | 0.81 | 1.06 | 0.96 | 0.99 | 1.1 | 0.94 | 0.98 |
| 8 | LCAT (hom) | 0.19 | 0.82 | 0.77 | 1.48 | 0.66 | 0.85 | 1 | 1 | 1 |
| 9 | DHCR7 | NA | NA | 0.2 | 1.13 | 0.37 | 0.54 | 1.13 | 0.37 | 0.54 |
| 10 | CYP7A1 | 0.97 | 2.09 | 1.74 | 0.93 | 1.21 | 1.15 | 0.98 | 1.04 | 1.03 |

Mutation numeric ID, gene, HDL, LDL and total cholesterol, from wet lab experiments[6], from predictions based on $f_{mut}$ as reported by van de Pas and colleagues[6], or trained $f_{mut}$ .

The effect of damaging mutations on the other genes of the test set have been simulated by reducing different set of rates of the model. The model predicted the effect of ABCA1 mutations as a decrease in HDL levels, but also produced overestimated decrease in LDL levels (Table 4), which is not usually observed in patients affected by related disease like Hypoalphalipoproteinemia [3]. CETP is a protein involved in the transport of cholesterol esters from HDL to LDL, deficiency of this protein can cause a marked increase of HDL levels [3]. In this case the model correctly predicted an increase in HDL cholesterol levels, with a bigger error when optimized *f_mut_* was used (Table 4). The LCAT gene is involved in cholesterol esterification in HDL particles, patients with mutations on this gene generally have low levels of HDL and LDL cholesterol [3]. In this case the model was not able to accurately simulate HDL and LDL levels in all cases (Table 3). Explanation could be that it was not possible to train the parameter for patients with a homozygous mutation on the LCAT gene (*f_mut_* has been assumed to be equal to 1). DHCR7 gene is involved in cholesterol biosynthesis pathway, mutations reducing related enzymatic activity cause low levels of blood cholesterol [12]. In all cases the model predicted a bigger decrease in total cholesterol levels with an error of −171.5%. CYP7A1 gene is involved in cholesterol catabolism and bile acids synthesis, mutations affecting this gene can cause an increase of total and hepatic cholesterol [13]. In this case the model generally predicted an increase of LDL and total cholesterol levels and a decrease in HDL cholesterol. Nevertheless, CYP7A1 simulations showed an underestimation of LDL and total cholesterol levels (Table 4).

### Performance assessment on the overall dataset

The overall assessment highlighted that the training phase increased model performance (Table 6). Both Pearson and Kendall correlation coefficients show that the use of trained *f_mut_* increased algorithm capability to predict variations on cholesterol levels caused by gene mutations. In particular, HDL levels predicted by the former version of the model have shown negative correlation with experimental values. The RMSE index computed on HDL and total cholesterol levels has been decreased thanks to the training procedure, and the same index on predicted LDL is one half of the one obtained with the original version of the model. R^2^ indices on blood cholesterol levels show an increase in the amount of variability explained by the model when a training phase is added.

**Table 6.** Models performances on the whole blind set.

| Prediction | PCC | KCC | RMSD | $R^2$ |
| --- | --- | --- | --- | --- |
| van de Pas et al predicted HDL ratio | -0.22 | -0.18 | 0.54 | 0.05 |
| genotype predicted HDL ratio | 0.32 | 0 | 0.4 | 0.11 |
| van de Pas et al predicted LDL ratio | 0.65 | 0.55 | 1.03 | 0.43 |
| genotype predicted LDL ratio | 0.74 | 0.5 | 0.5 | 0.55 |
| van de Pas et al predicted TC ratio | 0.66 | 0.63 | 0.71 | 0.43 |
| genotype predicted TC ratio | 0.75 | 0.69 | 0.57 | 0.56 |

Cholesterol level and predictor, Pearson Correlation Coefficient, Kendall rank Correlation Coefficient and R squared index computed on the test set

## Conclusions

In this work we improved and assessed the performance of an *in silico* prediction method for blood cholesterol levels. The addition of a training phase has generally improved model performance, as shown in Table 6. Our training phase overcomes the problem of model usability when no experimental data is available for *f_mut_* parameters estimation. The reducing parameters presented by van de Pas and colleagues were computed from variables obtained in wet lab experiments[6]. This procedure, in contrast with the training methodology we applied in this work, did not take in account that decreasing different rates by the same factor can lead to modification of cholesterol profiles with different magnitude. To better understand model responses to different simulations, we performed a sensitivity analysis on the rates involved in the test set (Fig 3). This analysis showed that reduction of rate 5 and 7 produced a consistent decrease of model predicted HDL, while increasing LDL and total cholesterol. The training procedure had computed *f_mut_* on the basis of the difference between experimental levels and model response to the reduction of selected rates, as previously explained. This procedure avoid an overestimation of the effect of mutations on the LDLR and APOB genes, as observed when *f_mut_* based on experimental variables were used (Table 4). The use of trained parameters has decreased prediction error when model was not able to correctly simulate the effect of a mutated gene on cholesterol levels. In particular rate 9 regulates the flow of cholesterol from free to esterified form in HDL particles, LCAT gene product activity. The effect of a mutation on this gene is predicted by the model as an increase of HDL cholesterol while the opposite is observed in real data (table 5). In this case the training procedures had hampered in part model inability to correctly predict HDL and LDL deviations caused by mutations on this gene by fixing the *f_mut_* to 1 in the homozygous case, since the reduction of this parameter was not able to reduce the difference between experimental and predicted values.

**Figure 3.**
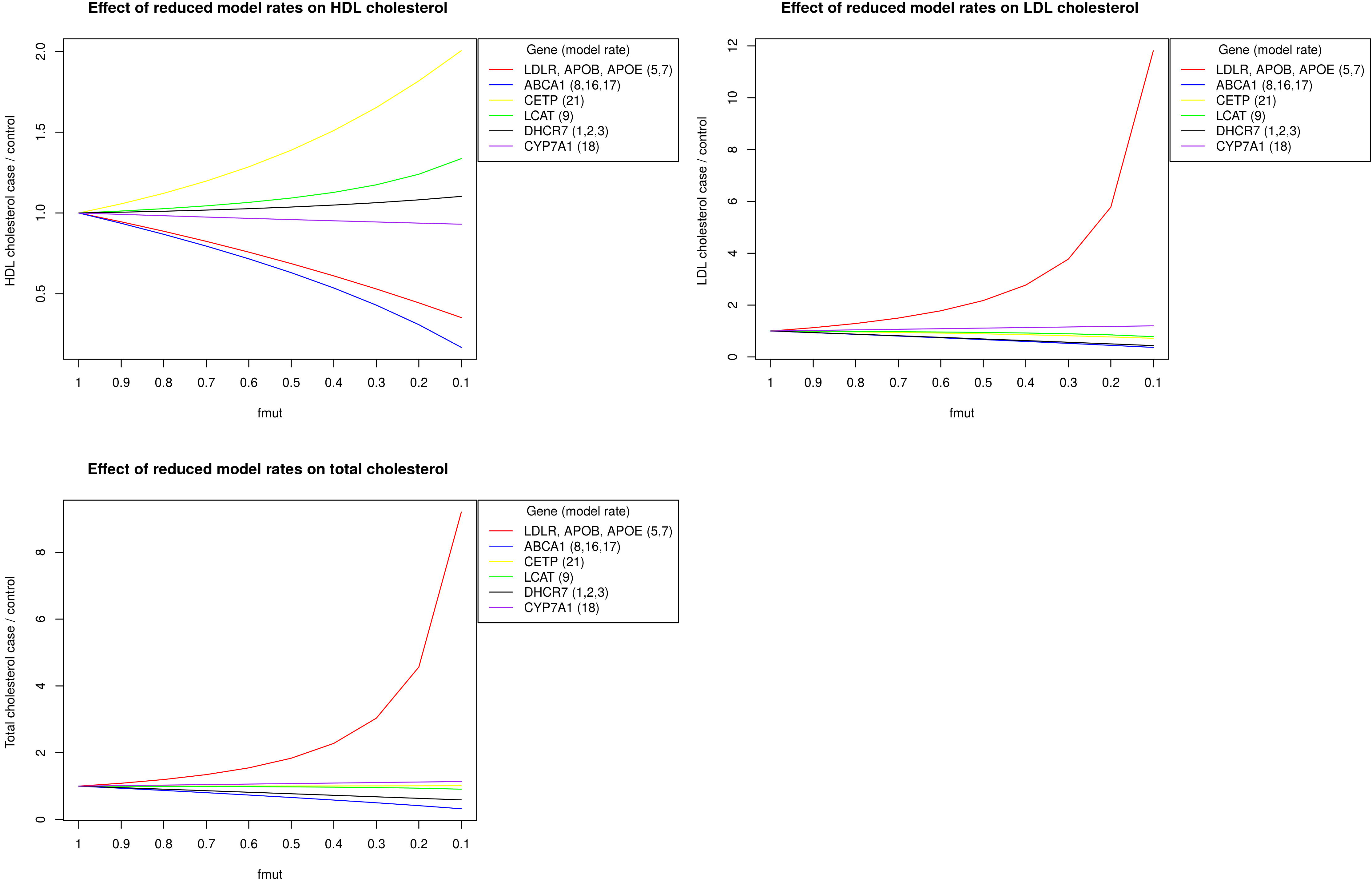
Model response in terms of HDL, LDL and total blood cholesterol at different values of *f_mut_*. The effect of reducing model rates, involved in the test procedure, on HDL, LDL or total cholesterol levels

This model can be considered a valid tool for the study of cholesterol metabolism in silico, considering the other models currently available [5] and the predictions error: the average relative deviations between model predictions and experimental data were 49% for HDL-C, 43% for LDL-C and 36% for total cholesterol [6]. Mathematical models are a simplified representation of the original system, this from one hand results in a relative simple tool for making inference and simulate different experimental conditions in silico. From the other hand, they don’t represent the selected system completely, hence deviation from real data are expected. Prediction error in principle could be decreased by increasing the number of parameters, however this process will increase model complexity and present problems related to parameter identifiability and fitting to experimental error [16]. Prediction of in silico cholesterol levels is a complex procedure, the physiologically based in silico cholesterol model optimized in this review has proven its ability to predict cholesterol levels behavior with reduced error when only genotype data is available. Given the huge number of genomic loci controlling cholesterol homeostasis, much of that still unknown, the effect of sex and environmental factors that affects blood cholesterol levels, the possibility of developing a software able to accurately predict cholesterol levels seems far from true. Hence we can consider the physiologically based in silico model of human cholesterol metabolism, optimized in this work, as a useful tool for studying the effect of damaging mutation on genes involved in cholesterol homeostasis

## Acknowledgments

The project was supported by the Italian Ministry of Health grant GR-2011-02346845. The authors are grateful to Francesco Tabaro for his useful suggestions. Ggplot2 has been used to produce Fig. 2[17]

